# The Arabidopsis PPR protein EMB1006 interacts with both EMB1270 and CFM2 to facilitate plastid *clpP1* intron2 splicing

**DOI:** 10.1101/2023.12.07.570680

**Authors:** Liqun Zhang, Yawen Zhu, Keyi Yang, Fangsheng Liao, Jiaqi Wang, Shuya Zhou, Jirong Huang, Yong-Lan Cui, Weihua Huang

## Abstract

Plastid intron splicing is an essential step for gene expression. Although many nucleus-encoded splicing factors have been identified for plastid intron splicing, how these proteins coordinately regulate intron splicing is unclear. Here we found that EMB1006, an Arabidopsis P-type PPR protein, is required for splicing several plastid introns including *clpP1* intron 2 by analyzing the level of intron retention in its knockdown lines and RIP-qPCR (RNA immunoprecipitation and quantitative PCR) assay. In addition, RNA electrophoretic mobility shift assay (REMSA) showed that EMB1006 specifically binds to the sequence UUACCAAACGU close to the 3’ end of *clpP1* exon 2 *in vitro*. Furthermore, yeast two hybrid (Y2H), split luciferase complementation (Split-LUC) and semi-*in vivo* pull-down assays showed that EMB1006 interacts with both EMB1270 and CFM2, which also interact each other. Taken together, our data indicate that EMB1006 forms a complex with EMB1270 and CFM2 to facilitate *clpP1* intron 2 splicing in chloroplasts.

## Introduction

Plastids and mitochondria are semi-autonomous organelles, where gene expression is highly regulated by numerous nucleus encoded proteins (de Longevialle et al., 2010). Pentatricopeptide repeat (PPR) proteins are one of the largest protein families in land plants and specifically regulate organellar gene expression at the post-transcriptional level. PPR proteins are defined as tandem arrays of the approximately 35-amino-acid long repeat (PPR motif), and each PPR protein contains 2 to 30 PPR repeats. They can be divided into P and PLS subfamilies, according to the characteristics of PPR motifs (Small, 2004). Each PPR protein binds to its specific RNA sequence based on the mechanism called one-repeat recognition of one-nucleotide. It has been well documented that PPR proteins play a wide range of roles in organellar RNA processing, such as stability, editing, cleavage, and intron splicing (Barkan and Small, 2014).

In the Arabidopsis plastid genome, there are twenty introns that are classified into group I (only one intron) and group II (19 introns), according to the splicing mechanisms (Huang et al., 2018; Zhang et al., 2021). Since group II introns have completely lost their self-splicing ability, a number of nucleus-encoded proteins are required for plastid intron splicing. Besides above mentioned PPR proteins (Wang et al., 2022), other RNA-binding proteins, such as Chloroplast RNA Splicing 1 (CRS1), CRS2-associated factors 1 (CAF1), CAF2, chloroplast RNA splicing and ribosome maturation (CRM) Family Member 2 (CFM2), CFM3, CFM1, RNC1 and WTF1, are also involved in group II intron splicing (Ostheimer et al., 2003; Asakura and Barkan, 2007; Barkan et al., 2007; Asakura et al., 2008; Feiz et al, 2021; Watkins et al., 2007; Kroeger et al., 2009). Interestingly, many of them contain the CRM domain, such as CRS1, CAF1, CAF2, CFM1, CFM2 and CFM3. RNC1, a ribonuclease III domain protein, was originally identified from the CAF1 and CAF2 coimmunoprecipitates in maize chloroplasts and played a role in a number of group II introns that include but are not limited to CAF1 and CAF2-dependent introns (Watkins et al., 2007). WTF1 (What’s This Factor 1) bearing the RNA-binding DUF860 domain was identified as another component of maize chloroplast RNPs and promoted group II intron splicing via interacting with RNC1 (Kroeger et al., 2009). RNC1 and WTF1 have been suggested to act as general splicing factors since their mutations disrupt the splicing of most plastid introns. In contrast, PPR proteins which bind to their single-stranded RNA targets in a modular and base-specific fashion can recognize specific sequences, and function at an initial sequence-specific binding step that alters RNA conformation ready for the recruitment of the relevant general splicing complex, such as the WTF1/RNC1 complex or CRM proteins (de Longevialle et al., 2010). Indeed, this hypothesis is supported by the recent studies from mitochondria and chloroplasts (Wang et al., 2020a; Zhang et al., 2021; Cao et al., 2022). For instance, PPR protein PPR14 is required for splicing of mitochondrial *NADH dehydrogenase 2* (*nad2*) intron 3 and *nad7* introns 1 and 2 through physical interaction with PPR-SMR1 and CRM protein Zm-mCSF1 in maize mitochondria (Wang et al., 2020a). Small PPR protein 2 (SPR2) containing merely four PPR repeats is involved in the splicing of more than half of the introns in maize mitochondria via interacting with small MutS-related domain protein PPR-SMR1 (Cao et al., 2022). Furthermore, it is found that SPR2 and/or PPR-SMR1 interact with other splicing factors, including PPR proteins EMPTY PERICARP16, PPR14, and CRM protein Zm-mCSF1, which participate in the splicing of specific intron(s) of the 13 introns. It suggested that SPR2/PPR-SMR1 serves as the core component of a splicing complex and possibly exerts the splicing function through a dynamic interaction with specific substrate recognizing PPR proteins in mitochondria (Cao et al., 2022). Consistently, in Arabidopsis chloroplasts, the PPR protein EMB1270 was shown to interact with CFM2 to splice specific group II introns (Zhang et al., 2021). These results imply that the combinations of organellar splicing factors contribute to specific group II intron splicing, in which PPR proteins play a sequence-recognition role and recruit other CRM proteins or other general splicing factors.

The caseinolytic protease (Clp) plays a central role in plastid development and function through selective removal of miss-folded, aggregated, or other unwanted proteins (Nishimura and van Wijk, 2015). The Clp system consists of a tetradecameric proteolytic core with catalytically active ClpP and inactive ClpR subunits, hexameric ATP-dependent chaperones (ClpC and ClpD) and adaptor protein(s) (ClpS1) enhancing delivery of subsets of substrates (Nishimura and van Wijk, 2015). Among these subunits, only ClpP1 is encoded by the chloroplast genome, while the rest are nucleus-encoded. Interestingly, *clpP1* contains no introns in monocots, but two introns in dicots (Kroeger et al., 2009; de Longevialle et al., 2010). It is reported that the splicing of Arabidopsis *clpP*1 intron 2 requires the three PPR proteins EMB1270/ACM1, EMB1006/ECD2, EMB976/PDM4, and a CRM domain protein CFM2 (Zhang et al., 2021; Wang et al., 2020b; Wang et al., 2021;Chen et al., 2020;). However, the mechanism by which these proteins collaborate to regulate *clpP*1 intron 2 splicing remains unclear.

Here we report that Arabidopsis PPR protein EMB1006 is complexed with EMB1270 and CFM2 to remove *clpP1* intron 2. We first predicted and verified the RNA sequence targeted by EMB1006 in *clpP1* exon 2. In addition, we found that EMB1006 interacts directly with EMB1270 and CFM2 *in vivo* and *in vitro*. Based on our previously reported results, EMB1270 binds to *clpP1* intron 2 and can also interact with CFM2, we proposed that the splicing of *clpP1* intron 2 requires both EMB1270 and EMB1006 proteins which bind to the different site of its pre-mRNA and then recruit a CRM domain RNA binding protein to form a splicing complex. Our findings provide new insights into how multiple PPR proteins regulate splicing of the same intron in plastids.

## Results

### Characterization of *emb1006* knockout mutant and the *EMB1006* co-suppression lines

The Arabidopsis EMB1006 (AT5G50280) with 11 PPR motifs belongs to the P-type subfamily, and its knockout mutant displayed the embryo lethal phenotype stopped at a globular stage (Meinke, 2020). We ordered a T-DNA insertion line CS16006 from the Arabidopsis Biological Resource Center (Figure 1A), and could not identify the homozygous mutant (*emb1006-1*). Careful examination of the siliques of heterozygous *emb1006-1/+* plants revealed that approximately one-fourth of the seeds were albino and aborted, confirming the embryo-lethal phenotype of *emb1006-1* (Figure 1B). We then transformed the *EMB1006* genomic DNA fused with 4×Myc-tag (*EMB1006-Myc*) at its C-terminal driven by its native promoter (a 1,745 bp region upstream of the start codon) into the heterozygous *emb1006-1/+* plants (Figure 1A). Complementation experiment showed that the *emb1006-1* homozygote plants expressing EMB1006-Myc fusion protein displayed the wild-type phenotype (Figure 1C, 1D and 1E), suggesting that EMB1006-Myc complements the embryo-lethal phenotype of *emb1006-1*.

**Figure 1.**
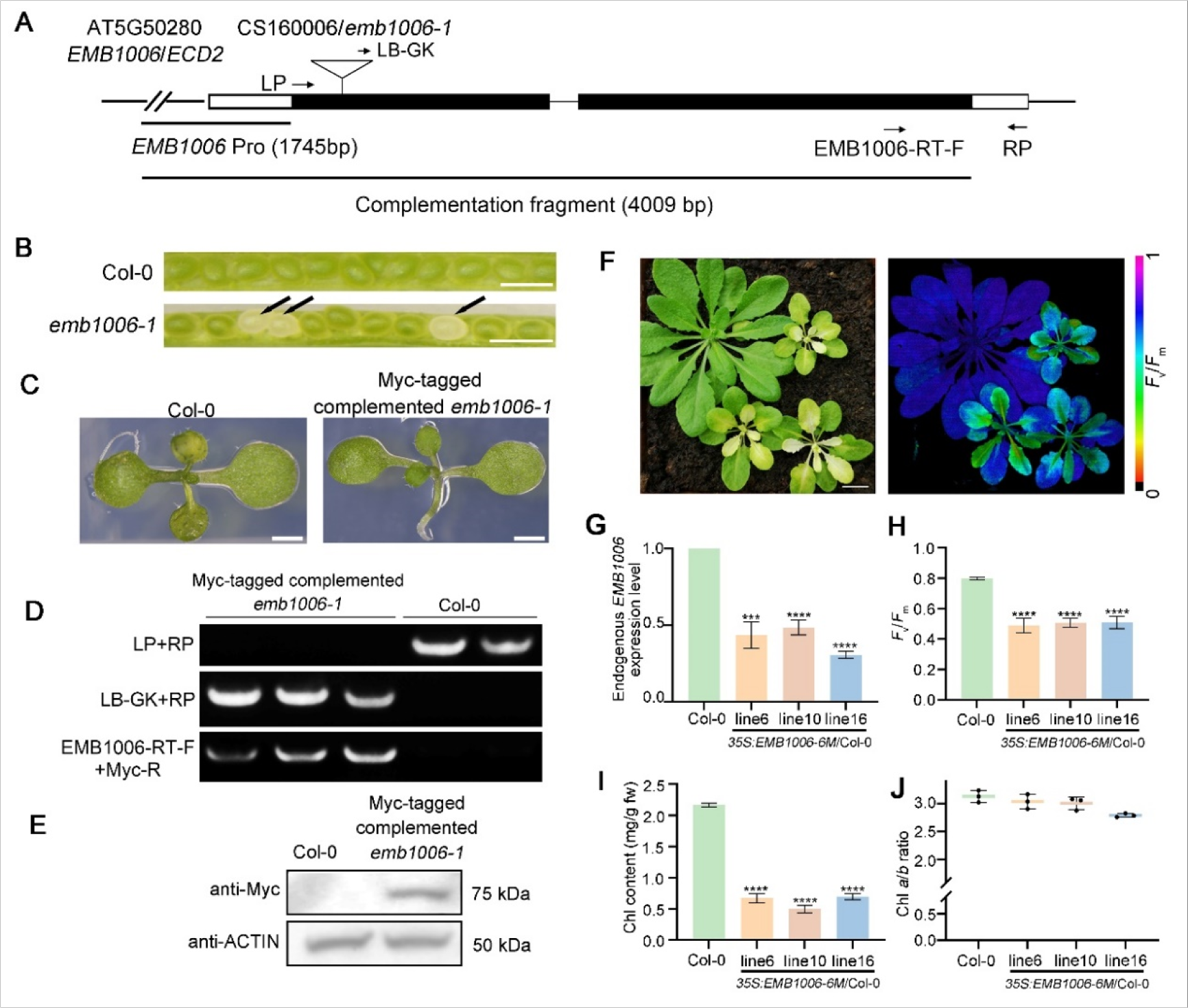
Characterization of *EMB1006* knockout mutant and knockdown lines. (A) Schematic diagram of *EMB1006* gene and the position of the T-DNA insertions. Black boxes, exons; white boxes, 5′- and 3′-untranslated regions; black lines, intron. The position of the T–DNA insertion in *emb1006-1* (CS160006) is indicated by the inverted triangle. The complementation fragment fused with Myc-tag is indicated by the straight line. The positions of the primers used for genotyping are indicated by the arrows. (B) Seed segregation in siliques from an *EMB1006*/*emb1006-1* heterozygous plant. Black arrows indicated the albino seeds. Scale bar = 1 mm. (C) Phenotypes of 10-day-old wild type and complemented *emb1006-1* seedlings. Scale bars = 1 mm. (D) Genotyping of Myc-tagged complemented *emb1006-1* seedlings by PCR. (E) Immunodetection of EMB1006-Myc protein in complemented *emb1006-1* plants. ACTIN was used as loading control. (F) Phenotypes (left panel) and *F*_v_/*F*_m_ values (right panel) of *EMB1006-6M* co-suppression lines. Scale bar = 15 mm. (G-J) Quantification of endogenous *EMB1006* expression levels (G), *F*_v_/*F*_m_ values (H), chlorophyll content (I), and chlorophyll *a*/*b* ratios (J) in wild type and *EMB1006* co-suppression lines. Data are mean ± SD (n=3). Asterisks indicate significant differences between wild type and *EMB1006* co-suppression lines (Student’s *t*-test, \*\*\**P* < 0.001 and \*\*\*\**P* < 0.0001).

To analyze the biochemical function of EMB1006, we knocked down *EMB1006* expression by overexpressing its truncated CDS fragment with the length of 1,506 bp from ATG (containing the first six PPR motifs) in the Col-0 background (Figure S1A), as previously reported (Chen et al., 2020). We found that about 20% of transgenic lines (*35S:EMB1006-6M*/Col-0) exhibited chlorosis and small rosette leaves at the early growth stage (Figure 1F), and about 65% of *35S:EMB1006-6M*/Col-0 plants had yellow stems and cauline leaves at the bolting stage (Figure S2A and S2B). The co-suppression phenotype is similar to that of *SOT5* (Chen et al., 2020). Three typical *35S:EMB1006-6M/*Col-0 co-suppression lines were chosen for further analysis. The expression level of the endogenous *EMB1006* gene significantly decreased in these co-suppression lines, compared with that in Col-0, confirming that the endogenous *EMB1006* is partially silenced (Figure 1G). The chlorosis rosette leaves of *35S:EMB1006-6M/*Col-0 plants had a significantly lower *F*_v_/*F*_m_ value and total chlorophyll content than those of Col-0 (Figure 1F, 1H and 1I). However, the ratio of chlorophyll *a* to *b* did not significantly alter in these co-suppression lines, compared to that in Col-0 (Figure 1J).

### *EMB1006* Encodes a Chloroplast-Localized PPR Protein

EMB1006 is predicted to target chloroplasts and has a transit peptide of 63 amino acids (https://services.healthtech.dtu.dk/services/TargetP-2.0/). To confirm the subcellular localization of EMB1006, we overexpressed EMB1006 fused with YFP at the C-terminal (EMB1006-YFP) or the N-terminal 123 amino acids of EMB1006 fused with YFP (EMB1006_1-123_-YFP) in Arabidopsis protoplasts. Confocal microscopy analysis showed that the YFP fluorescence was co-localized with chlorophyll autofluorescence, suggesting that EMB1006 is a plastid-targeted protein (Figure 2A).

**Figure 2.**
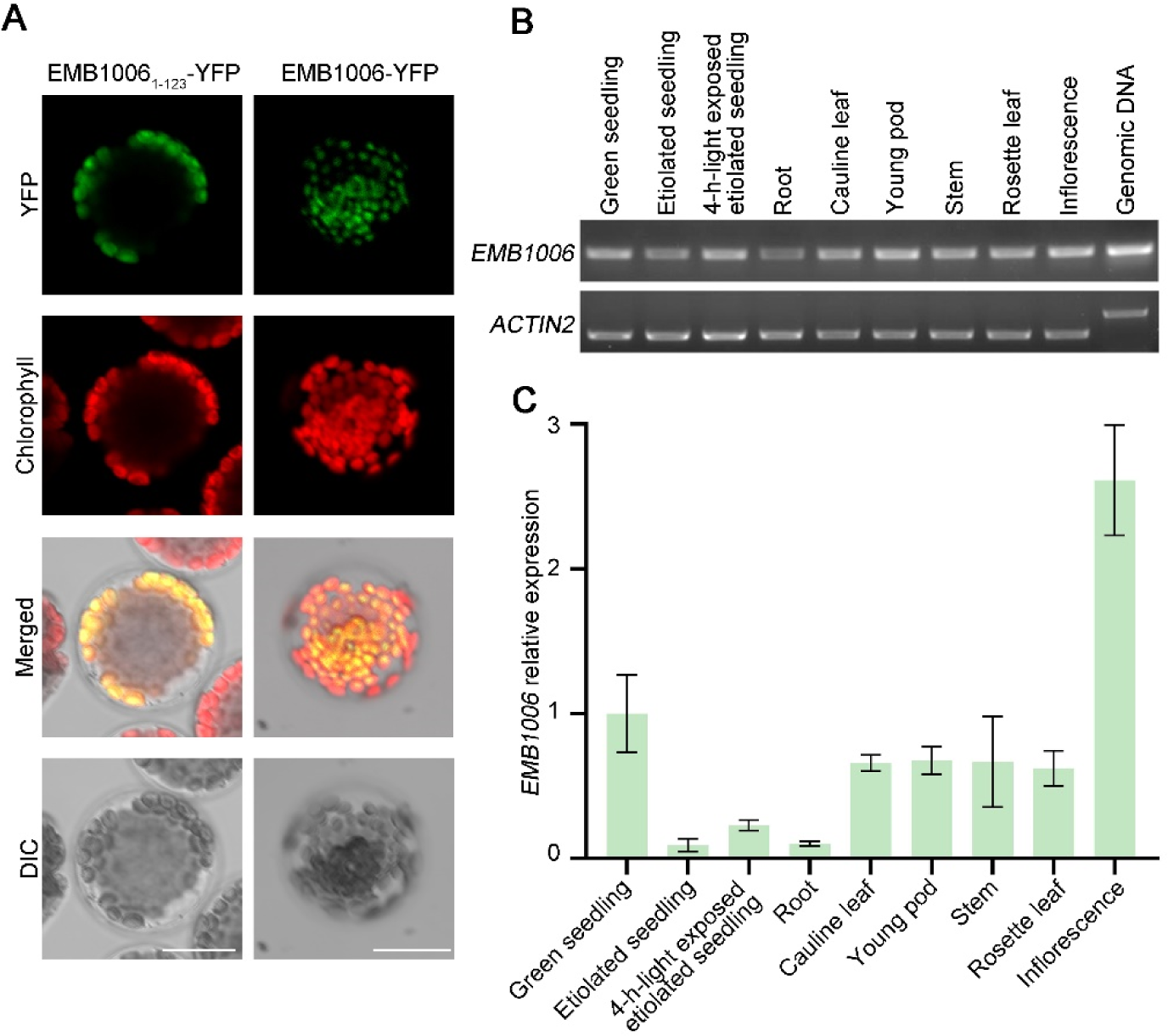
Subcellular localization of EMB1006 protein and its gene expression pattern. (A) Microscopy analysis of the YFP fusion protein EMB1006_1-123_-YFP and EMB1006-YFP transiently in Arabidopsis protoplasts. The green fluorescence of EMB1006_1-123_-YFP and EMB1006-YFP was overlapped with chloroplast autofluorescence in merged images. DIC, Differential interference contrast microscopy. Scale bar = 15 μm. (B and C) RT-PCR(B) and RT-qPCR(C) analyses of EMB1006 expression in various seedlings and tissues. Genomic DNA was used as a template control, and *ACTIN2* expression was used as a loading control. The RT-qPCR data are means ± SD (n=3).

The eFP Browser (http://bar.utoronto.ca/efp_arabidopsis/cgi-bin/efpWeb.cgi) database showed that *EMB1006* is ubiquitously expressed in almost all the tissues and organs. Then we performed semi-quantitative RT-PCR and RT-qPCR to detect the expression pattern of *EMB1006*. The results showed that *EMB1006* was highly expressed in green tissues including leaves, siliques, stems, and inflorescences, but mildly expressed in etiolated seedlings and roots (Figure 2B, 2C).

### EMB1006 is required for intron splicing of several plastid genes

Considering that P-type PPR proteins are usually involved in organelle RNA splicing, we analyzed the effect of *EMB1006* knockdown on plastid gene splicing. Semi-quantitative RT-PCR analysis showed that *rps12* intron 2 and *clpP1* intron 2 strongly retained in the *35S:EMB1006-6M*/Col-0 lines, compared with those in Col-0 (Figure 3A). RT-qPCR analysis confirmed that the splicing efficiency of *rps12* intron 2 and *clpP* intron 2 was dramatically reduced in *35S:EMB1006-6M*/Col-0 lines. And the splicing efficiency of *ndhA* intron and *ycf3* intron1 was mildly reduced (Figure 3B). These results were consistent with the report that the splicing defects of plastid *rps12* intron 2, *clpP1* intron 2, *ycf3* intron 1 and *ndhA* intron in *EMB1006*/*ECD2* RNAi lines (Wang et al., 2021). To confirm whether EMB1006 specifically binds to *clpP1*, *rps12*, *ycf3* and *ndhA* transcripts, we performed RIP assays using complementation lines expressing the EMB1006-Myc fusion protein in the *emb1006-1* background. Expression of the fusion protein EMB1006-Myc was verified by immunoblotting with a monoclonal antibody against Myc (Figure 3C). RIP-qPCR assays showed that *clpP1*, *rps12*, *ycf3* and *ndhA* transcripts but not the *rpl2*, *LHCB1* and *ACTIN2* transcripts were enriched in the immunoprecipitated fraction from EMB1006-Myc plants, compared with the control only expressing Myc (Col-Myc) (Figure 3D). These results indicate that EMB1006 associates with the plastid *clpP1* intron 2, *rps12* intron 2, *ycf3* intron 1 and *ndhA* intron *in vivo*.

**Figure 3.**
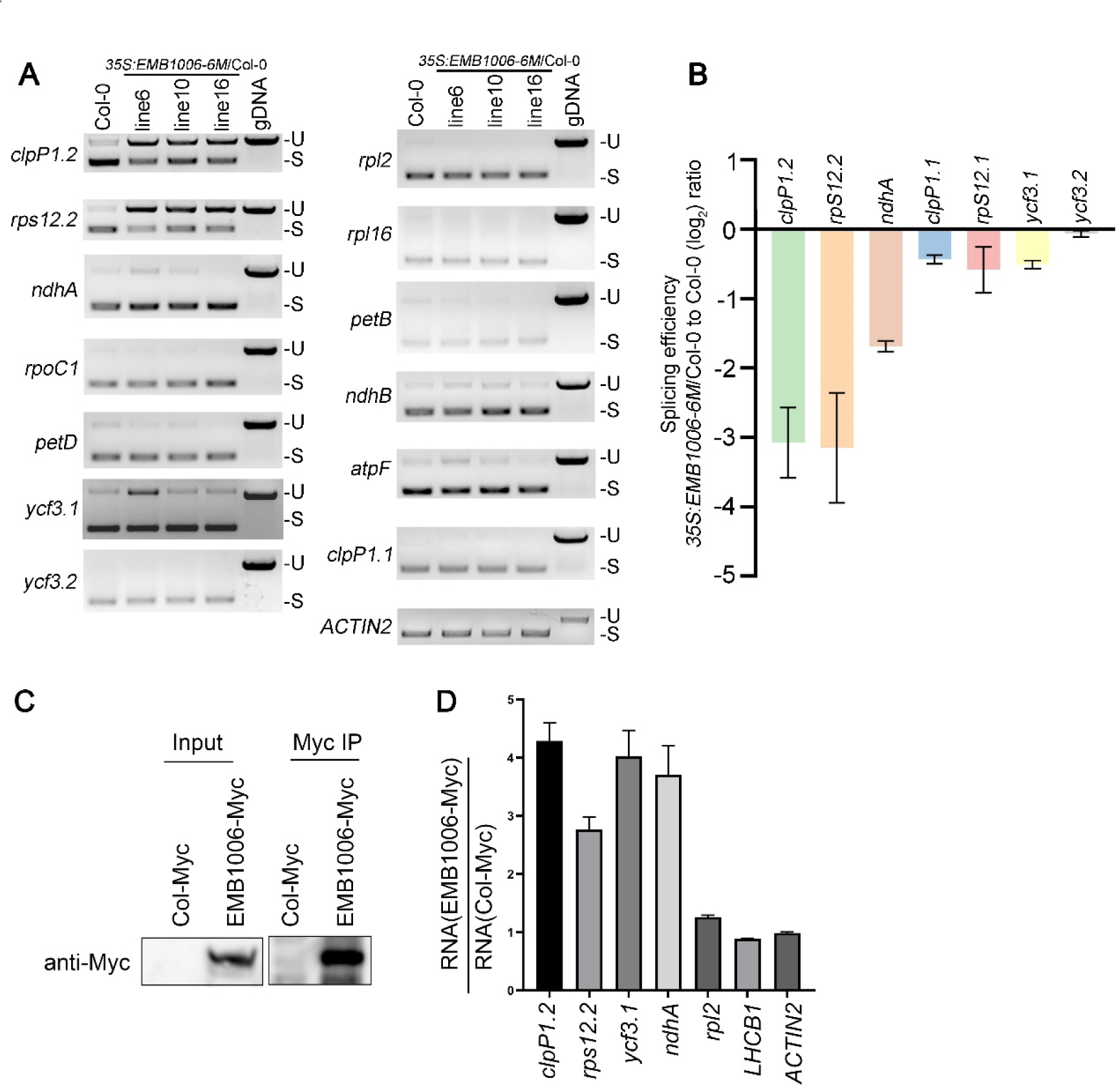
Splicing defects in plastid introns in the *EMB1006* co-suppression lines and EMB1006 associate with its targets RNA in *vivo*. (A) Semi-quantitative RT-PCR analysis of chloroplast introns splicing in the *EMB1006* co-suppression lines. Genomic DNA (gDNA) was used as a template control, and *ACTIN2* expression was used as a loading control. Three biological replicates were analyzed. S, spliced; U, unspliced. (B) RT-qPCR analysis of the splicing efficiency of seven chloroplast mRNA introns in the *EMB1006* co-suppression lines. Data are mean ± SD (n=3). (C) Immunoblots of proteins present in crude leaf extracts from EMB1006-Myc and Col-Myc transgenic plants and RNA immunoprecipitated using anti-Myc antibody. (E) RT-PCR analysis of *clpP1.2, rps12.2, ycf3.1* and *ndhA* introns present in crude leaf extracts from EMB1006-Myc and Col-Myc transgenic plants. Plastid *rpl2*, nuclear encoded genes *LHCB1* and *ACTIN2* transcripts were used as a negative control. Bars represent ± SD of three biological replicates.

### EMB1006 is required for accumulation of ClpP1 and plastid ribosomal proteins

The *35S:EMB1006-6M*/Col-0 co-suppression lines exhibit chlorosis in leaves and stems, implying that photosynthetic machinery cannot accumulate in these knockdown lines. Based on the above result showing that EMB1006 is particularly required for splicing of *clpP1* and *rps12* introns, we examined their protein expression levels in co-suppression lines. Western blot assay confirmed that the abundance of ClpP1 strongly decreased in *35S:EMB1006-6M*/Col-0 lines, compared with that in Col, whereas the mild reduction of other chloroplast proteins, such as ClpC1, VAR2 and LHCB1, were also detected between *35S:EMB1006-6M*/Col-0 and Col-0 plants (Figure 4A and 4B). In addition, the plastid encoded AtpB was also strongly reduced in the co-suppression lines. Since it has been demonstrated that the reduced level of ClpP protease is usually accompanied with a lowered accumulation of mature plastid rRNA (Wu et al, 2014; Zhang et al, 2021), we predicted that the whole ribosome complex would be reduced in the *35S:EMB1006-6M*/Col-0 lines. Because the plastid RPS12 antibody was not available, we tried to detect some other plastid ribosome subunits in the co-suppression lines. The results showed that the plastid ribosomal protein RPS2 was dramatically decreased and RPL10, RPL16, and RPS1 were mildly reduced in the co-suppression lines (Figure 4A and 4B). These results suggested that EMB1006 is required for accumulation of ClpP1, plastid ribosomal proteins and other chloroplast proteins.

**Figure 4.**
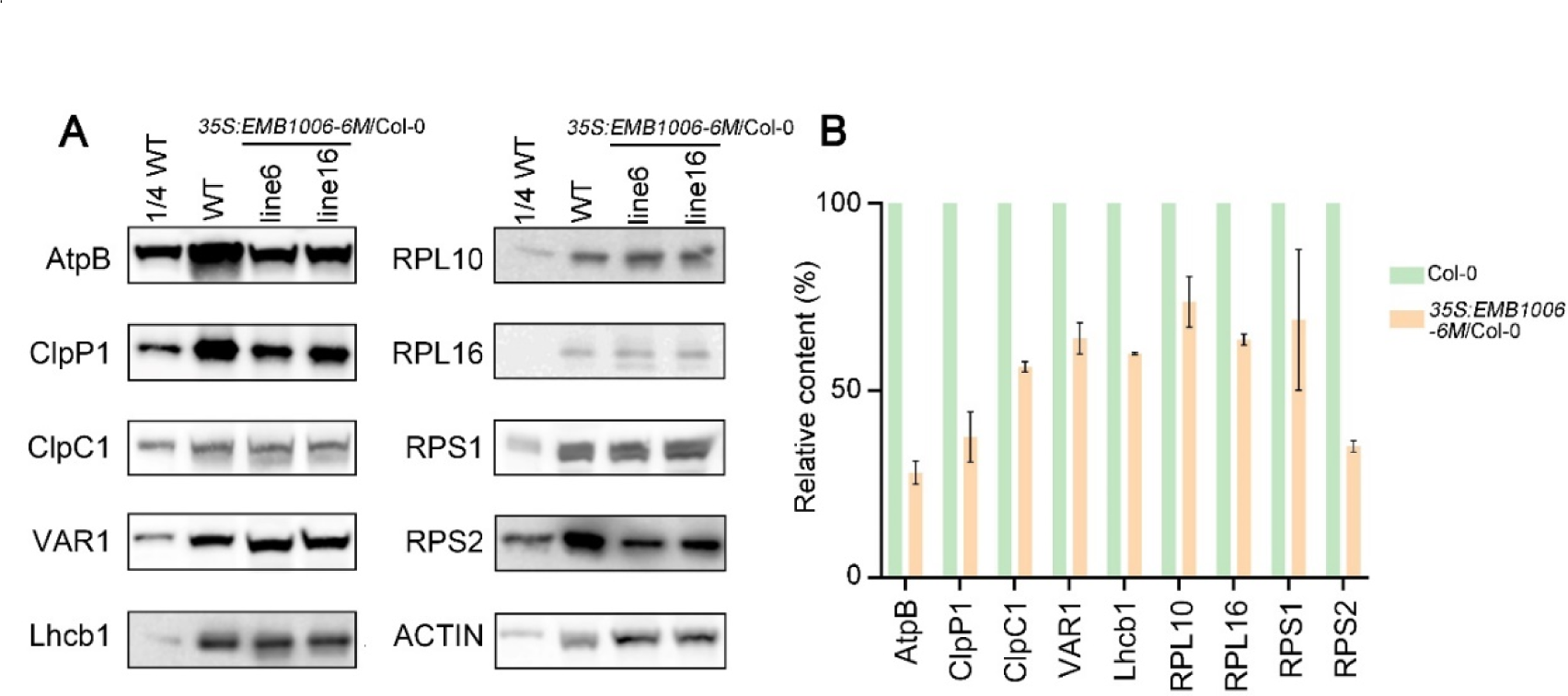
Reduced ClpP1 and plastid ribosomal proteins in the *EMB1006* co-suppression lines. (A) Immunoblot analysis of chloroplast proteins in *EMB1006* co-suppression lines. ACTIN was used as loading control. (B) Proteins bands in (A) were analyzed by ImageJ software. Values (means ± SD; n = 2) are given as ratios to protein amounts in the *EMB1006* co-suppression lines and wild type.

### EMB1006 directly binds to *clpP* exon 2

ScanProsite (https://prosite.expasy.org/) analysis showed that EMB1006 has 11 P-type PPR motifs at the C terminal and no domains at the N terminal. To identify the substrates for EMB1006, we predicted the binding sequence of EMB1006 to be BNMHYRRRHBG (B = U/C/G, D = A/G/U, N = A/U/C/G, K = G/U, M = A/C, H = A/C/U, Y = C/U, R = A/G) via the PPRCODE web server (http://yinlab.hzau.edu.cn/pprcode) (Yan et al., 2019). Interestingly, one of the predicted EMB1006-binding candidates is present in the 3’ end of *clpP1* exon 2 (Figure 5A and B). To verify the prediction, we expressed and purified MBP-fused EMB1006 from *E. coli* (Figure 5C). REMSA assays showed that EMB1006 but not the MBP tag bound to the probe Cy5-*clpP1-*exon 2 (CY5: AAUUACCAAACGUAUAGCAUUCC) (Figure 5D). In addition, the binding of EMB1006 to the probe was blocked with an increase in the concentration of the non-labeled probe, but not by the non-labeled *clpP1-*intron 2 probe (Figure 5E). It is worth noting that the non-labeled *clpP1* intron 2 probe was the target of EMB1270 (Zhang et al, 2021). These results indicate that EMB1006 and EMB1270 facilitate *clpP1* intron 2 splicing through their specific binding to the target sequence.

**Figure 5.**
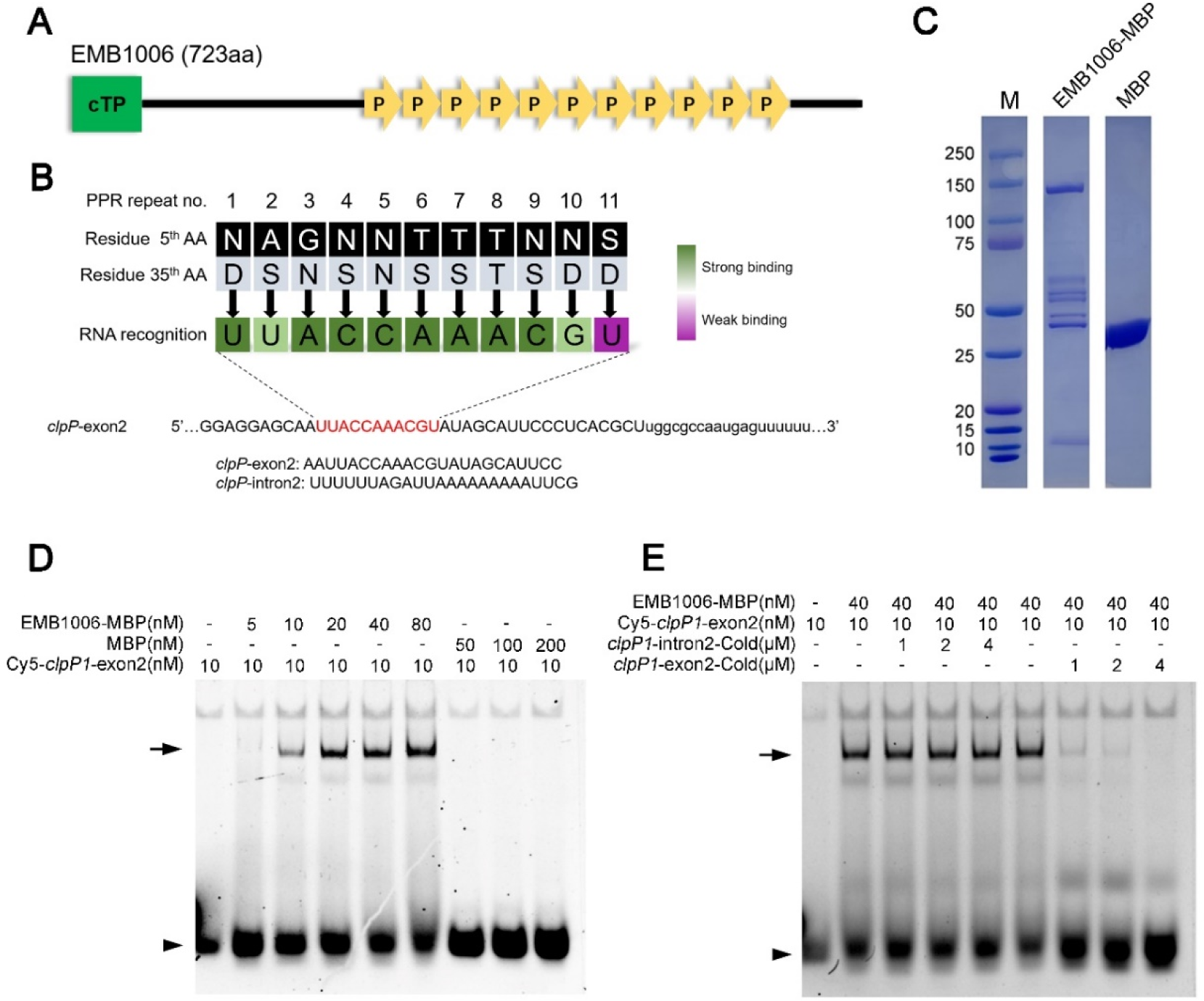
RNA electrophoretic mobility shift (REMSA) assay showed that EMB1006 directly binds to *clpP1* exon 2. (A) Schematic diagram of the EMB1006 protein containing 11 PPR motifs. The recombinant EMB1006 protein without transit peptide was used for the REMSA. (B) The predicted 11 nucleotides targeted by the 11 PPR motifs of EMB1006. In each motif, the two key amino acid residues and their nucleotide targets are shown. The coloring of the motif–nucleotide pairs indicate predicted binding strength. (C) The purified recombinant EMB1006-MBP (middle lane) and MBP (right lane) were separated by SDS-PAGE and visualized by Coomassie Brilliant Blue (CBB) staining. (D) REMSA assay demonstrated that EMB1006-MBP binds to the *clpP1* exon2 probe. The concentrations of probes, recombinant EMB1006, and MBP are indicated above each lane. (E) Competition experiment using non-labeled *clpP1* exon2 and *clpP1* intron2 probe. Arrow and arrowheads indicate the EMB1006-MBP-RNA complex and free labeled RNA, respectively. Arrows indicate the EMB1006-MB-RNA complex and arrowheads indicated the free probes.

The splicing defect of *clpP1* intron 2 was also reported in the co-suppression lines of *EMB976,* which encodes a P-type PPR protein with 22 PPR motifs (Chen et al, 2020). We tried to find a possible target sequence of EMB976 with 21 nt in *clpP1* intron2 (Table S1) as well as to expressed and purified a recombinant EMB976. However, REMSA assay showed that the recombinant EMB976 failed to bind the labeled probe (Figure S2A and 2B). Interestingly, Wang et al. (2020) reported that the knockout mutant of *EMB976*/*PDM4* had a defect in the splicing of *clpP1* intron 1 but not *clpP1* intron 2 besides *ndhA*, *petB*, *ycf3* intron1, *petD* intron. These results indicated that *clpP1* intron 2 probably is not the direct target of EMB976. Thus, the true target of EMB976 needs to be clarified.

### EMB1006 directly interacts with the plastid splicing factors CFM2 and EMB1270 *in vitro* and *in vivo*

The above results showed that Arabidopsis EMB1006 is required at least for the splicing of chloroplast *clpP1* intron 2. To date, several plastid splicing factors, such as CFM2, EMB1270 and EMB976, have been reported to target the same plastid genes as EMB1006 (Asakura et al, 2007; Zhang et al, 2021; Chen et al, 2020), while the WTF1-RNC1 splicing complex and MatK are associated with the splicing of plastid *rps12* intron2 (Kroeger et al, 2009). We proposed that these splicing proteins may interact with each other to regulate intron splicing in plastids. To test this hypothesis, we examined whether EMB1006 interacts with CFM2, EMB1270, EMB976, MatK or WTF1. Y2H analysis showed that EMB1006 interacted with CFM2 and EMB1270 (Figure 6B), but not with WTF1, MatK and EMB976 (Figure 6C). In addition, EMB976 did not interact with CFM2 (Figure 6C). Consistently, co-expression analysis showed that *CFM2 vs EMB1006*, *CFM2 vs EMB1270*, and *EMB1006 vs 1270* gene pairs are highly co-expressed (Figure S3), which indicates they may form a protein complex to conduct intron splicing in plastid.

**Figure 6.**
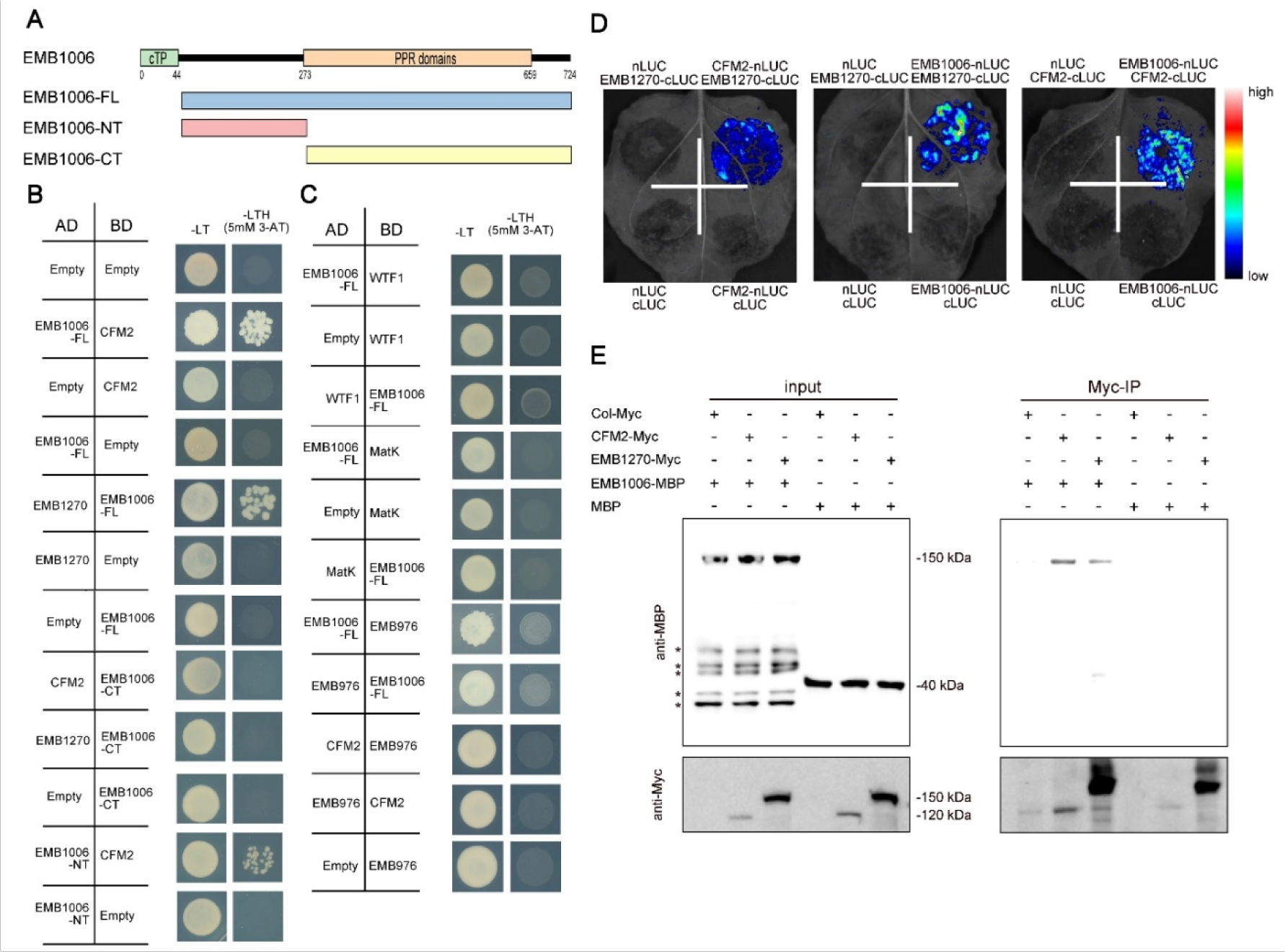
EMB1006 interacts with EMB1270 and CFM2. (A) Schematic diagrams of EMB1006 proteins used in Y2H. C-terminal EMB1006 and N-terminal EMB1006 were constructed based on the presence or absence of the PPR structural domain. EMB1006-FL indictes full-length EMB1006. EMB1006-NT indictes EMB1006 amino acids 44-273. EMB1006-CT indictes EMB1006 amino acids 273-724, which include 11 PPR motifs. (B) EMB1006 interacts with the CFM2 and EMB1270, rspectively in the Y2H assays. (C) EMB1006 do not interact with WTF1, MatK, EMB976 in the Y2H assays. And EMB976 do not interact with CFM2 in the Y2H assays. The transformed yeast cells were grown on -LT (SD-Leu/-Trp) mediums and -LTH+3-AT (SD-Leu/-Trp/-His with 5 mM 3-AT) mediums. (D) Split-LUC assays indicating the interaction of EMB1006 with EMB1270 and CFM2 in tobacco (*Nicotiana benthamiana*) leaves. The interaction between EMB1270 and CFM2 acts as a positive control. (E) Semi-*in vivo* pull-down assays showed the interactions of EMB1006-MBP with EMB1270-Myc or CFM2-Myc. The total protein extracts from *EMB1270:EMB1270-Myc*/*emb1270* and *CFM2:CFM2-Myc*/*cfm2* plants served as bait, and purified EMB1006-MBP *in-vitro* served as preys. The bait proteins and prey protein pulled down by EMB1270-Myc and CFM2-Myc were detected with anti-MBP and anti-Myc antibodies, respectively. Asterisks indicate the non-specifically band.

To map the region of EMB1006 that interacts with CFM2 or EMB1270, we divided EMB1006 into the N-terminal (Non-PPR domain) and C-terminal (11 PPR motifs) parts (Figure 6A). Our results showed that CFM2 interacted with the N-terminal (EMB1006-NT), whereas EMB1270 cannot physically interact with the C-terminal (EMB1006-CT) (Figure 6B), suggesting that the Non-PPR domain of EMB1006 interacts with EMB1270.

To detect the above interaction *in vivo*, we performed split luciferase complementation (Split-LUC) assays in *Nicotiana benthamiana* leaves. Our results showed that high levels of luciferase activity were detected in the combination of co-expressing by co-expressing CFM2 fused to N-terminal luciferase (CFM2-nLUC) and EMB1270 fused to C-terminal luciferase (EMB1270-cLUC), which is consistent with our previous report that EMB1270 interacted with CFM2 *in vivo* by Co-IP (Zhang et al, 2021). Similarly, high levels of luciferase activity were detected in the combination of co-expressing EMB1006 fused to N-terminal luciferase (EMB1006-nLUC) and EMB1270 fused to C-terminal luciferase (EMB1270-cLUC) (Figure 6D), indicating the interaction between EMB1006 and EMB1270. Likewise, high levels of luciferase activity were also detected in the combination of co-expressing EMB1006-nLUC) and CFM2-cLUC (Figure 6D), indicating of the interaction between EMB1006 and CFM2.

We further analyzed the interaction between EMB1006 and CFM2 or EMB1270 through a semi-*in vivo* pull-down assay. The total protein was extracted from the complementation transgenic plants expressing the Myc-tagged CFM2 (*CFM2-Myc*/*cfm2*) and EMB1270 (*EMB1270-Myc*/*emb1270*), respectively. While the recombinant protein MBP-EMB1006 and MBP were purified from *E. coli*. Immunoprecipitation analysis showed that CFM2-Myc or EMB1270-Myc interacted with MBP-EMB1006 but not with MBP (Figure 6E). Taken together, these results suggested that EMB1006 directly interacts with CFM2 and EMB1270.

## Discussion

### EMB1270, EMB1006 and CFM2 bind to the different sites of *clpP1* pre-mRNA and probably form a RNP complex to facilitate *clpP1* intron2 splicing

In this study, we found that Arabidopsis EMB1006 interacts with EMB1270 and CFM2 *in vitro and in vivo*. Combined with our previous report that EMB1270 and CFM2 interact each other (Zhang et al., 2021), it is likely that EMB1006, EMB1270 and CFM2 are in the same protein complex to facilitate intron splicing. In addition, RIP-qPCR and REMSA assays showed that EMB1006 targets the specific sequence close to the 3’ end of *clpP1* exon 2, while EMB1270 targets the sequence in the *clpP1* intron 2 (Figure 7A and 7B). Based on these results, a working model was proposed for Arabidopsis plastid *clpP1* intron 2 splicing (Figure 7C). First, PPR protein EMB1006 and EMB1270 bind to their respective sites of *clpP1* pre-mRNA and change the RNA conformation. These two PPR protein interacted with each other and then recruit CRM domain protein CFM2 which can also bind *clpP1* pre-mRNA. The suitable protein-RNA complex was finally formed to help the intron to be spliced (Figure 7C). Given that there are a large number of PPRs in land plants and some of them share the same targets. Probably it is general that the different sites of group II intron were recognized by different PPRs, which then recruit other general splicing factors such as CRM domain protein to change pre-mRNA conformation that susceptible to be spliced. Consistently, a splicing complex composed of different PPRs and CRM domain proteins for maize mitochondrial intron splicing was reported (Cao et al, 2022). The mechanism is similar to those PPR proteins involved in RNA editing in chloroplasts and mitochondria. Numerous PPR proteins can combine with universal editing complex proteins (MOF/RIP, ORRM, PPO1, and OZ1) and their family members to form specific editing complexes for editing specific sites (Andres Colas et al., 2017; Yan et al., 2017; Sun et al., 2016; Wang, et al, 2023)

**Figure 7.**
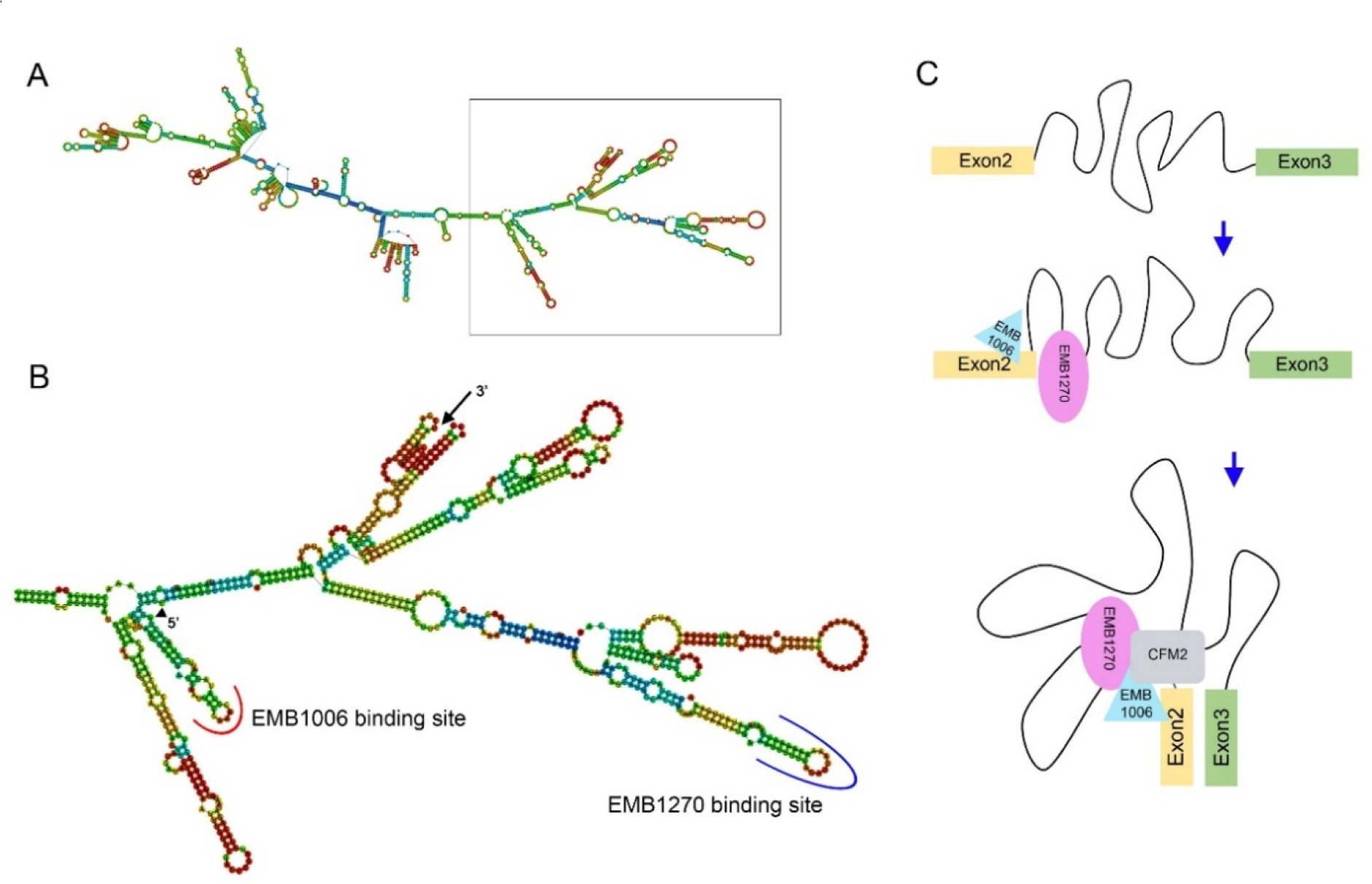
Binding site of EMB1006 and EMB1270 in *clpP1* mRNA and a proposed working model for Arabidopsis plastid *clpP1* intron2 splicing. (A) The secondary structure of *clpP1* pre-mRNA predicted by RNAfold WebServer (http://rna.tbi.univie.ac.at/cgi-bin/RNAWebSuite/RNAfold.cgi). (B) Magnification of the rectangle in (A) showing the the sequence of *clpP1* intron2. Arrowhead indicate the 5’ of *clpP1* intron2. Arrow indicate the 3’ of *clpP1* intron2. The red line indicates the binding site of EMB1006 and the blue line indicates the binding site of EMB1270. (C) A proposed working model for Arabidopsis plastid *clpP1* intron2 splicing process involved with EMB1006, EMB1270 and CFM2.

### EMB976 is probably involved in *clpP1* intron2 splicing indirectly

Although EMB976 is reported to be required for *clpP1* intron 2 splicing, however, in this study, it is showed the recombinant EMB976 cannot bind the predicted sequence found in *clpP1* intron2 *in vitro.* And no interactions were detected between EMB976 and EMB1006, EMB976 and CFM2 by Y2H and split-LUC assay (data not show). Considering *clpP1* mRNA cannot be immune-precipitated from EMB976/PDM4-GFP complementary seedlings, although *clpP1* intron1 splicing is defective in the *pdm4* mutant (Wang, et al, 2020), we suppose *clpP1* intron2 is probably not the direct target of EMB976.

## Materials and Methods

### Plant materials and growth conditions

The Arabidopsis ecotype Columbia-0 (Col-0) was used as the wild type (WT) in this study. The complementary plants *EMB1270:EMB1270-Myc*/*emb1270* and *CFM2:CFM2-Myc*/*cfm2* were previously described (Zhang et al., 2021). The *emb1006-1* (CS160006) mutant was obtained from ABRC stock center (https://abrc.osu.edu/) and were genotyped by PCR using gene specific primers (Table S3). Co-suppression lines of *35S:EMB1006-6M*/Col-0 was identified from T_1_ generation. The lines with typical co-suppression phenotypes were selected for physiological experiments in the T_2_ generation. Seeds were surface-sterilized by 75% ethanol, 0.05% Triton and stratified at 4℃ for 3 days, and then sown onto half-strength Murashige and Skoog (MS) medium with 0.5% sucrose. Seeds were also directly sown in soil and grown in a phytotron with long-day conditions (16 h light/8 h dark) and light intensity (100 μmol photons m^-2^ s^-1^) at 22℃.

### Complementation of *emb1006-1* mutant

For mutant complementation, the 4,009-bp wild type genomic fragment containing the putative EMB1006 promoter was amplified by Fastpfu (TransGen), cloned into the pCAMBIA1300 vector modified with four tandem Myc tags using *pEASY*®-Basic Seamless Cloning and Assembly Kit (TransGen Biotech). The resulting construct was transformed into *Agrobacterium tumefaciens* GV3101 competent cells and introduced into heterozygous *emb1006-1* plants by the floral dip method. The complemented homozygous plants were identified among the obtained transgenic plants by PCR and Western-blot analysis.

### Analyses of Chl content and Chl fluorescence

20 mg of leaves tissue collected from 2-week-old *Arabidopsis* plants was ground to a fine powder in liquid nitrogen. Total Chl was extracted with the buffer (0.2 M Tris-HCl, pH 8.0: acetone: H_2_O = 1:8:1) for 5 min vortex in the dark. The levels of Chl *a* and Chl *b* were determined as our previously described (Huang et al., 2018). Chl fluorescence was analyzed using Imaging PAM 101 (Walz), and *F*_v_/*F*_m_ values were determined according to the manufacturer’s instructions.

### Subcellular localization of EMB1006_1-123_ and EMB1006 fused with YFP

To express YFP-fused protein, the 369 bp at the 5’ end of *EMB1006* CDS and the full length genomic of *EMB1006* gene without stop code were amplified from the cDNA of wild type seedlings using the primers (Table S3). The sequences were cloned into the pENTR SD/D-TOPO entry vector (Invitrogen) and then recombined into the p2GWY7 vector (http://gateway.psb.ugent.be/vector/show/p2GWY7/search/index/). The purified plasmid was transformed into the *Arabidopsis* protoplasts according to the method (Wu et al., 2009).

### RNA extraction, RT-PCR and RT-qPCR

Total RNA was extracted from plants using RNA Easy Fast Plant Tissue Kit (Tiangen Biotech). First-strand cDNA was synthesized with a PrimeScript™ RT reagent kit with gDNA Eraser (Takara). For gene expression pattern analysis, total RNA was extracted from rosette and cauline leaves, stems, inflorescences, siliques, and roots of 40-d-old plants, and 5-d-old seedlings grown in light or darkness or etiolated seedlings exposed to light for 4 h, respectively. RT-PCR and RT-qPCR analyses of chloroplast RNA splicing were conducted using the chloroplast gene-specific primers (Table S3). The data were normalized using *ACTIN2* as a reference.

### Protein expression, purification and RNA electrophoretic mobility shift assay (REMSA)

The CDS of *EMB1006* and *EMB976* was synthesized by GenScript, and the codon were optimized with the GenSmart^TM^ Codon Optimization tool for higher expression in the prokaryotic expression system. The optimized CDS was cloned into expression vector pMAL-c5x. Protein expression in *E. coli* strain Rosetta was induced by treatment with 0.2 mM isopropyl-β-D-thiogalactopyranoside at 16°C for 16 h. The MBP-fused protein was purified with Amylose Resin (E8021V; New England Biolabs) according to the manufacturer’s protocol. The RNA probes were chemically synthesized and their 5′-ends labeled with Cy5 (Sangon Biotech). The sequences of the probes were as follows:

Cy5-*clpP1*-exon2: (CY5) AAUUACCAAACGUAUAGCAUUCC

*clpP1*-exon2-Cold: AAUUACCAAACGUAUAGCAUUCC

*clpP1*-intron2-Cold: UUUUUUAGAUUAAAAAAAAAUUCG

Cy5-*rps12*-exon2: (CY5) AAACCAAACUCUGCUUUACGU

*rps12*-exon2-Cold: AAACCAAACUCUGCUUUACGU

Cy5-*clpP1*-intron2-976: (CY5) GAAGAGGUUUUUCAAAUGAUA

For REMSA, the RNA probes and EMB1006/EMB976 protein were incubated at room temperature for 30 min in reaction solution (1 mM MgCl_2_, 10 mM HEPES, pH 7.3, 20 mM KCl, 5% glycerol (v/v), 0.1 μg tRNA, and 1 mM dithiothreitol). The reactants were analyzed by electrophoresis in a native polyacrylamide gel, and signals were detected using an Azure Biosystems C600 Imager.

### Protein Extraction and Immunoblot

Protein extraction and immunoblot analyses were performed as described in Huang et al. (2013). The ClpP1, ClpC1, AtpB and Lhcb1 antibodies were purchased from Agrisera Company (https://www.agrisera.com/). The following antibodies were purchased from PhytoAB Company: RPL10 (PhytoAB, PHY0423), RPL16 (PhytoAB, PHY0431A), RPS1 (PhytoAB, PHY0424A) and RPS2 (PhytoAB, PHY0427). They are used at a 1:1,000 dilutions. The protein bands were quantified by Image J software and normalized to Actin.

### RIP assay

1 g of seedlings collected from 2-week-old *Arabidopsis* plants on 1/2 MS medium was ground to a fine powder in liquid nitrogen. About 50 mg powder was taken out for RNA extraction as input. The samples were homogenized in 5 mL RIP lysis buffer (100 mM KCl, 2.5 mM MgCl_2_, 50 mM Tris-HCl, pH 7.4, 0.5 mM phenylmethylsulfonyl fluoride, 0.2% Nonidet P-40 (v/v), one-fourth of a complete proteinase inhibitor Cocktail tablet (Roche), and 50 units/mL RNase inhibitor (Sangon Biotech). Cell debris was removed by centrifugation for 5 min at 12,000 *g* at 4℃. The clarified lysates were incubated with 150 μL Myc magnetic beads (Biomake) for 4 h at 4℃. The magnetic beads were washed six times with RIP lysis buffer at 4°C and divided for RNA and protein analyses. The RNA was recovered by incubating the magnetic beads in 500 μL proteinase K buffer (300 mM NaCl, 10 mM EDTA, 0.1 M Tris-HCl, pH 7.4, 1 μg/μL proteinase K (Sangon Biotech), and 2% SDS (v/v)) for 15 min at 65°C. The RNA was extracted with phenol:chloroform and precipitated with 2.5 volumes of ethanol. cDNA was synthesized from the PrimeScript™ RT reagent Kit with gDNA Eraser (TaKaRa) for RT-qPCR.

### Yeast two hybrid assay

The CDS removal of transit peptide coding sequences of *EMB1006*, *EMB1006-NT*, *EMB1006-CT*, *CFM2*, *EMB1270*, *WTF1*, *MatK* and *EMB976* were cloned into the pENTR SD/D-TOPO entry vector (Invitrogen) and then recombined into pDEST22 and pDEST32 destination vectors, respectively. For yeast two-hybrid assays, plasmids were transformed into yeast strain AH109 by the lithium chloride–polyethylene glycol method according to the manufacturer’s manual (Weidi Company). The transformants were selected on SD-Leu-Trp plates. The protein–protein interactions were tested on SD-Trp-Leu-His plates with or without 5 mM 3-amino-1,2,4-triazole (3-AT).

### Split Luciferase Complementation Assays

The full length of EMB1006 gDNA, CFM2 and EMB1270 cDNA fragments without stop codes were cloned into the 936-pDEST-GW-NLUC and 937-pDEST-GW-CLUC vectors. These constructs were transformed into *Agrobacterium tumefaciens* GV3101 competent cells, respectively. Positive strains were shaken overnight in LB medium with kanamycin and rifampicin. The GV3101 was pellet and re-suspended in liquid MS, adjusting the OD_600_ to 0.6∼0.8. GV3101 cells harboring the indicated combinations of constructs expressing nLUC and cLUC fusion proteins were mixed at a ratio of 1:1 and were co-infiltrated into 4-week-old tobacco (*Nicotiana benthamiana*) leaves, which were kept in the dark for 2 d. The leaves were incubated with 1 mM D-luciferin sodium salt substrate (Yeasen, China) for 10 min before luciferase activity detection with a luminescence imaging workstation (Tanon 5200).

### Semi-*in vivo* Pull-down Assays

For EMB1006 interaction assays, the preys recombinant MBP-EMB1006 and MBP were purified from Rosetta strains of *E. coli* as previously described (Zhang et al., 2021). The bait CFM2-Myc and EMB1270-Myc crude chloroplast protein was extracted from *Pro-CFM2:CFM2-Myc*/*cfm2* seedlings and Pro-*EMB1270:EMB1270-Myc*/*emb1270* seedlings, respectively (Zhang et al., 2021), and solubilized with IP buffer (25 mM Tris-HCl, pH 7.5, 150 mM NaCl, 1 mM EDTA, 1% Triton X-100,10% glycerol) containing 1× Protease Inhibitor Cocktail (Roche) and incubated for 30 min at 4℃. The samples were centrifuged for 15 min at 13,000 g. After centrifugation, the supernatant, the prey recombinant protein, and 5 μL Myc antibody (Millipore) was incubated for 2h at 4℃. The magnetic Protein A/G Beads (Bimake, B23201) was then added to the mix and the mix was incubate overnight at 4℃. The magnetic beads were washed for 6 times with wash buffer (25 mM Tris-HCl, pH 7.5, 150 mM NaCl, 1 mM EDTA, 0.5% Triton X-100,10% glycerol). The MBP protein was added to the binding reaction as a negative control. The precipitates were eluted into 30μL 1×SDS loading buffer and detected using anti-Myc and anti-MBP antibody (1:5,000, New England Biolabs).

## Supplemental data

Figure S1. Generation of *EMB1006* co-suppression lines

Figure S2. Recombinant EMB976-MBP did not bind the Cy5-*clpP1*-intron2-976 probe, which was predicted to be the target of EMB976.

Figure S3. Co-expression analysis of *EMB1006*, *CFM2* and *EMB1270*.

Table S1. The position of the predicted target sequences of EMB1006, EMB1270 and EMB976 in *clpP1*.

Table S2. EMB1006 and EMB976 codon optimized sequence.

Table S3. Primers used in this study.

Table S4. Gene accession numbers.

## ACKNOWLEDGMENTS

We thank ABRC for providing the *emb1006-1* (CS160006) seeds. We thank Zhilei Mao, Tongtong Guo and Minqing Liu for protein interactions experiments. We thank Shui Wang for providing 936-pDEST-GW-NLUC and 937-pDEST-GW-CLUC vectors. This work was supported by the National Natural Science Foundation of China (32171293), The Fund of Innovation Program of Shanghai Municipal Education Commission (2021 - 01 - 07 - 00 - 02 - E00117), and Shanghai Engineering Research Center of Plant Germplasm Resources (17DZ2252700).

## AUTHOR CONTRIBUTIONS

W.H., L.Z. Y.C. and J.H. designed the experiments; L.Z., Y.Z., K.Y., J.W. F.L., S.Z., and W.H. performed the experiments. W.H., L.Z, Y.C.and J.H. analyzed the data and wrote the manuscript. All authors read and approved of this manuscript.

